# Fast corrective responses in redundant motor control are shaped by intrinsic constraints of movement patterns

**DOI:** 10.64898/2026.03.11.711221

**Authors:** Toshiki Kobayashi, Daichi Nozaki

**Author notes:** Corresponding authors: Toshiki Kobayashi & Daichi Nozaki.

## Abstract

In real-world motor tasks, body movements unfold in a high-dimensional space, whereas task errors are defined in a lower-dimensional space. How such low-dimensional errors propagate across redundant motor degrees of freedom to generate rapid corrective responses remains poorly understood. To address this question, we developed a redundant bimanual task in which participants manipulated a virtual stick with both hands to move its tip to a visual target. Visual perturbations either displaced the stick tip (end-effector relevant errors) or altered the tilt angle of the stick without affecting the tip position (end-effector irrelevant errors), allowing us to dissociate errors defined in task space from those arising in redundant dimensions. Participants rapidly corrected both types of perturbations using highly stereotyped movement patterns. Corrections to end-effector relevant errors consistently involved coordinated changes across redundant dimensions, and even perturbations that did not affect task success elicited systematic corrective responses. Together, these results demonstrate that fast corrective responses in redundant motor control are not generated by flexible re-optimization, but are instead shaped by intrinsic coordination constraints that govern how visual errors propagate through the motor system.

## Introduction

Accurate control of limb end-effectors, such as the hand, is essential for interacting with the external environment and relies on multiple error correction processes operating at different timescales. One such process is rapid feedback control, which allows ongoing movements to be corrected in response to unexpected perturbations with latencies faster than voluntary reactions (Scott, 2016). Another process involves updating motor plans based on past errors, enabling feedforward compensation in subsequent movements and promoting motor learning (Shadmehr & Mussa-Ivaldi, 1994; Thoroughman & Shadmehr, 1999; Takiyama et al., 2015). These error correction processes have been predominantly studied using unimanual reaching tasks that constrain movement to a two-dimensional plane and limit kinematic degrees of freedom to the shoulder and elbow (Morasso, 1981; Flash & Hogan, 1985; Gordon et al., 1994). Importantly, these paradigms involve relatively limited kinematic redundancy, as the dimensionality of the control variables matches that of the task space.

In contrast to these constrained paradigms, the motor system typically possesses more degrees of freedom than are strictly necessary to control an end-effector (Bernstein, 1996). In such redundant systems, multiple body configurations can produce the same end-effector outcome, giving rise to dimensions of movement that directly affect task success as well as dimensions that do not. A central challenge for motor control is therefore how task-relevant errors, defined in a low-dimensional task space, are distributed across high-dimensional motor degrees of freedom to achieve reliable control. One influential framework for addressing redundancy is optimal feedback control theory, which proposes that motor behavior follows a minimal intervention principle: variability is actively corrected in task-relevant dimensions while being tolerated in task-irrelevant dimensions (Todorov & Jordan, 2002; Todorov, 2004; Scott, 2004).

To investigate how redundancy is resolved during movement, we previously developed a redundant bimanual task in which participants manipulated the endpoint of a virtual stick using both hands (Kobayashi & Nozaki, 2024). Although the same endpoint position could be achieved through multiple stick configurations, participants consistently adopted stereotyped movement patterns involving stick tilt, indicating intrinsic coordination constraints linking movement dimensions that directly affect the control of the end-effector with those that do not. Based on this background, we defined the terms ‘end-effector relevant’ and ‘end-effector irrelevant’, thereby distinguishing them from the commonly used terms ‘task-relevant’ and ‘task-irrelevant’ (Kobayashi & Nozaki, 2024). The constraints between end-effector relevant and end-effector irrelevant movements appear to shape how adapted motor commands are organized during movement planning. However, it remains unknown how these coordination constraints influence the rapid propagation of end-effector relevant errors through redundant motor spaces during ongoing movement. The present study addresses this question by examining how the motor system rapidly reconfigures redundant effectors to correct end-effector errors via feedback control.

## Results

### Stereotyped movement patterns manipulating a stick with both hands

A total of 41 healthy right-handed adults participated in a bimanual stick-manipulation task. Participants held a virtual stick with both hands (length: 40 cm; inter-hand distance: 15 cm) and controlled the position of either the right tip (N = 21) or the left tip (N = 20) of the stick (Fig. 1a; Fig. S1a). Because the task was kinematically redundant, identical tip trajectories could, in principle, be produced by many different combinations of hand movements. For example, participants could translate the stick by moving both hands in parallel without changing its orientation, or they could achieve the same tip displacement by tilting the stick (Fig. 1b). During the baseline phase (280 trials), participants moved the tip from a starting point to a target located 15 cm away in one of seven directions (0°, ±30°, ±60°, ±90°). Despite the availability of multiple movement solutions, participants exhibited highly consistent movement patterns. The trajectories of the tip and both hands were approximately straight (Fig. 1c; Fig. S1b), and the hand closer to the controlled tip was consistently moved farther than the other hand, regardless of whether the left or right tip was controlled.

**Figure 1.**
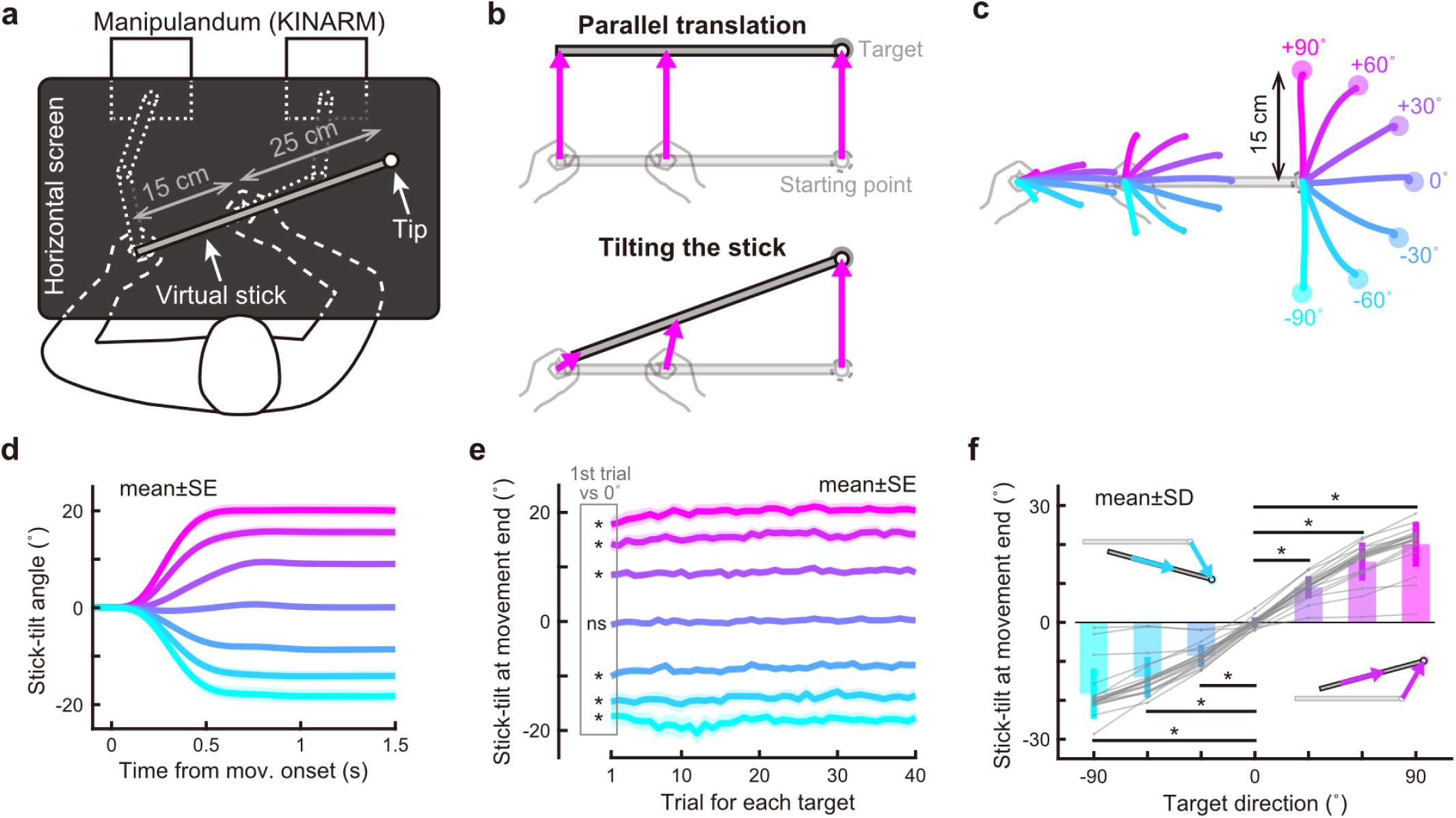
Kinematics to move the right tip of the stick with both hands. (**a**) Participants held handles with both hands and moved the tip of a virtual stick on the screen. (**b**) They can reach a visual target by moving both hands parallel (*top*) or tilting the stick (*bottom*). (**c**) During the baseline phase, participants were instructed to move the tip from a home position (a pale picture of the stick) to a target that appeared in one of seven directions. Movement trajectories of the left hand, right hand, and tip were drawn over the diagram (averaged across participants). (**d**) The time-course of the stick-tilt angle during the reaching movements. They achieved the task while tilting the stick. Each color represents the direction of the target, as shown in **c**. The shaded areas indicate SEM. (**e**) Trial-to-trial changes in the stick-tilt angle at the movement end. They adopted the strategy of tilting the stick from the beginning of the experiment (t-test compared with zero, |t(20)| > 8.8, Bonferroni corrected p < 0.001). The shaded areas indicate SEM. (**f**) The stick-tilt angle at the movement end significantly differs compared to that when aiming at a horizontal target (paired *t*-test, |t(20)| > 11.8, Bonferroni corrected p < 0.001). Gray plots indicate individual data. The error bars mean SD. *: p < 0.001.

Notably, participants rarely translated the stick by moving both hands in parallel, except when aiming at the 0° target. Instead, they systematically tilted the stick during movement execution, with stick tilt gradually increasing from movement onset (Fig. 1d; Fig. S1c). This tilting strategy was present from the very first trial and was significantly different from zero for all non-zero target directions (t-test compared with zero, |t(20)| > 8.8, Bonferroni corrected p < 0.001, Fig. 1e; |t(19)| > 10.9, Bonferroni corrected p < 0.001, Fig. S1d). The final stick-tilt angle was strongly modulated by target direction, increasing monotonically with the angular deviation of the target from the horizontal (paired t-test compared with the stick-tilt when aiming at the 0° target, |t(20)| > 11.8, Bonferroni corrected p < 0.001, Fig. 1f; |t(19)| > 13.3, Bonferroni corrected p < 0.001, Fig. S1e). Regardless of whether participants controlled the left or right tip, they consistently rotated the stick clockwise for clockwise targets and counterclockwise for counterclockwise targets (Fig. S1f). This directional tuning of stick tilt was highly stereotyped across participants and trials.

Thus, although the task allowed many equivalent solutions in terms of end-effector kinematics, participants consistently adopted a specific coordination pattern linking end-effector displacement to systematic changes in a end-effector irrelevant dimension. These stereotyped movement patterns indicate the presence of intrinsic coordination constraints between end-effector relevant and irrelevant dimensions. Consistent with previous work, this baseline coordination structure likely reflects minimization of overall hand movement distance (Kobayashi & Nozaki, 2024).

### Rapid corrective responses to end-effector perturbations

During the second half of the task, visual perturbations were introduced while participants reached a target located at 0°. Nineteen participants (right-tip control: N = 10; left-tip control: N = 9) performed perturbation trials in which the position of the stick was unexpectedly shifted by ±3 cm during movement execution (Fig. 2a; Fig. S2a). Perturbations were applied pseudo-randomly once every three trials to minimize predictive correction. Because movements toward the 0° target were typically performed without stick tilt during baseline, participants could, in principle, correct the perturbation by translating the stick in parallel (Fig. 2b). Alternatively, if feedback corrections were constrained by the baseline coordination structure, correcting the tip position error would necessarily involve systematic changes in stick tilt (Fig. 2c).

**Figure 2.**
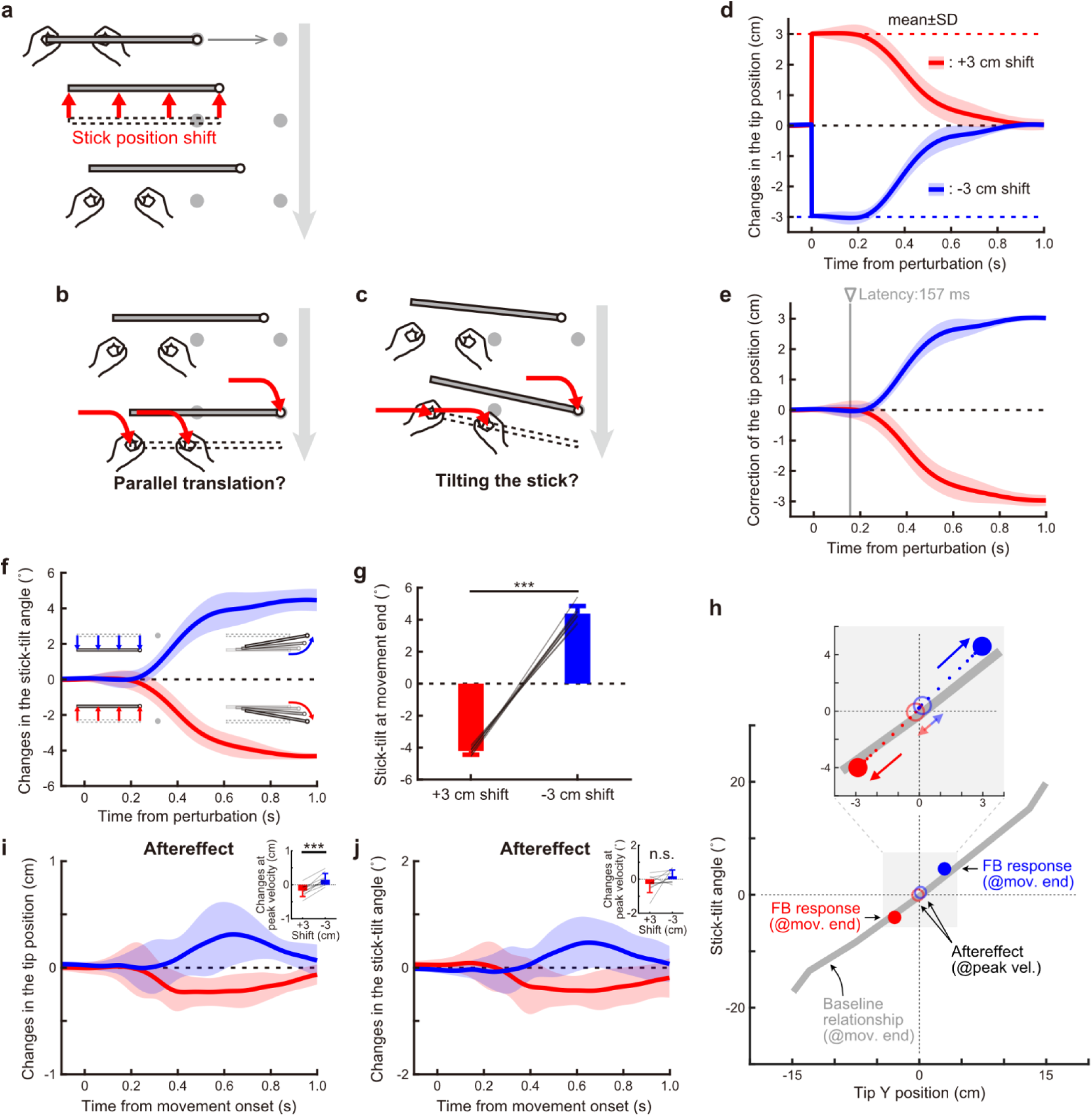
Corrective rapid response to the end-effector relevant perturbation. (**a**) During the perturbation phase, a cycle comprising two null trials and a visual perturbation trial was repeated. In the perturbation trial of a stick position shift, the stick was displaced by +3 or −3 cm orthogonally to the target direction when the tip was 1 cm away from the starting point. The stick-tilt angle kept the same as the tilting angle of both hands. (**b**) Participants could correct the tip error by translating both hands in parallel. (**c**) Alternatively, they could correct the error by tilting the stick. (**d**) The time-course changes in the tip position orthogonal to the target direction. The red (blue) plot shows average data when the stick was shifted by 3 cm (−3 cm). The shaded areas indicate the SD. (**e**) Changes in the correction amount of the tip position error. The participants rapidly corrected errors with a latency of 157 ms. Latency was derived from the time derivative of the position profiles (i.e., the velocity profiles). (**f**) Changes in the stick-tilt angle show that the correction of the tip position was accompanied by a stick rotation. Insets depict the stick on the monitor, exaggeratedly drawn for clarity. (**g**) Changes in the stick-tilt angle at the movement end show that the stick rotation significantly accompanied the correction of the tip position (paired *t*-test, t(9) = −39.972, p < 0.001). The error bars indicate the SD. (**h**) A plane representing the relationship between the tip position and the stick-tilt angle. The tip Y position denotes the Y-coordinate relative to the starting position. The bold gray line indicates the relationship in the baseline phase. The red (blue) filled circles and red (blue) open circles represent the feedback responses at the movement end and aftereffects at the peak velocity, respectively. The red (blue) small dots in the inset represent trajectories from perturbation onset to movement end at 50 ms intervals. All plots show data averaged across participants. (**i, j**) In trials following the perturbation trials, biases of the tip position and stick-tilt angle were observed as aftereffects of the perturbations (insets: paired t-test, tip position: t(9) = −5.781, p < 0.001, stick-tilt angle: t(9) = −2.032, p = 0.073). ***: p < 0.001.

Participants initiated corrective responses to the tip displacement with a latency of approximately 157 ms (which was derived from the velocity profiles) (Figs. 2d, 2e), consistent with rapid visuomotor feedback control. Importantly, these corrective responses were accompanied by systematic rotations of the stick. Upward corrections of the tip position were consistently associated with counterclockwise stick rotation, whereas downward corrections were accompanied by clockwise rotation (Fig. 2f). These feedback-induced changes in stick tilt closely matched the direction-dependent coordination patterns observed during unperturbed baseline movements (Fig. 1f). Thus, although the perturbation affected only the end-effector position, corrective responses propagated into the redundant stick-tilt dimension in a highly stereotyped manner. At the end of the movement, the stick remained significantly tilted in the direction induced during the corrective response (Fig. 2g; paired t-test, t(9) = −39.972, p < 0.001). Similar patterns were observed when participants controlled the left tip of the stick (Figs. S2b–d), indicating that the observed coordination was not specific to the controlled side.

The results demonstrate that rapid corrective responses to end-effector relevant perturbations are shaped by intrinsic coordination constraints and thus are accompanied by end-effector irrelevant errors. To further examine this speculation, we performed an additional analysis to test whether the feedback response pattern followed the relationship between tip movement and stick angle in the baseline phase. The result showed that the tip position and stick-tilt angle during the feedback response were consistent with the baseline relationship (Fig. 2h).

An important question is how such corrective responses influence subsequent motor learning. Specifically, with respect to the secondary deviations which were resulted from feedback correction to a perturbation, does the motor system attempt to compensate for the deviations in the following trial, or are they left uncorrected? To address this question, we examined trial-by-trial aftereffects following perturbation trials. The trials following a perturbation were normal trials without any visual perturbation. Participants showed aftereffects biased the initial tip position downward (upward), consistent with predictive compensation for the perturbation (Fig. 2i; paired t-test, t(9) = - 5.781, p < 0.001). Additionally, they showed systematic aftereffects of the stick angle that matched the direction of the corrective response expressed in the preceding trial, rather than counteracting it, although the effects were not significant (Fig. 2j; paired t-test, t(9) = −2.032, p = 0.073). Similar to the feedback response, these movement biases followed the baseline relationship (Fig. 2h).

Finally, we examined whether visual information about stick orientation was necessary for these corrective responses by introducing a perturbation in which the stick disappeared except for the controlled tip (Fig. 3a; Fig. S3a). Despite the absence of visual information about stick orientation, feedback corrections exhibited the same systematic coupling between tip correction and stick tilt as observed when the full stick was visible (Figs. 3b–d; Figs. S3b–d). Moreover, aftereffects in both tip position and stick angle were indistinguishable from those observed in the standard condition (Figs. 3e, f; paired t-test, tip position: t(9) = −2.701, p = 0.024, stick-tilt angle: t(9) = −1.604, p = 0.143). These results indicate that the propagation of end-effector errors into stick-tilt changes does not depend on explicit visual feedback about the end-effector irrelevant dimension.

**Figure 3.**
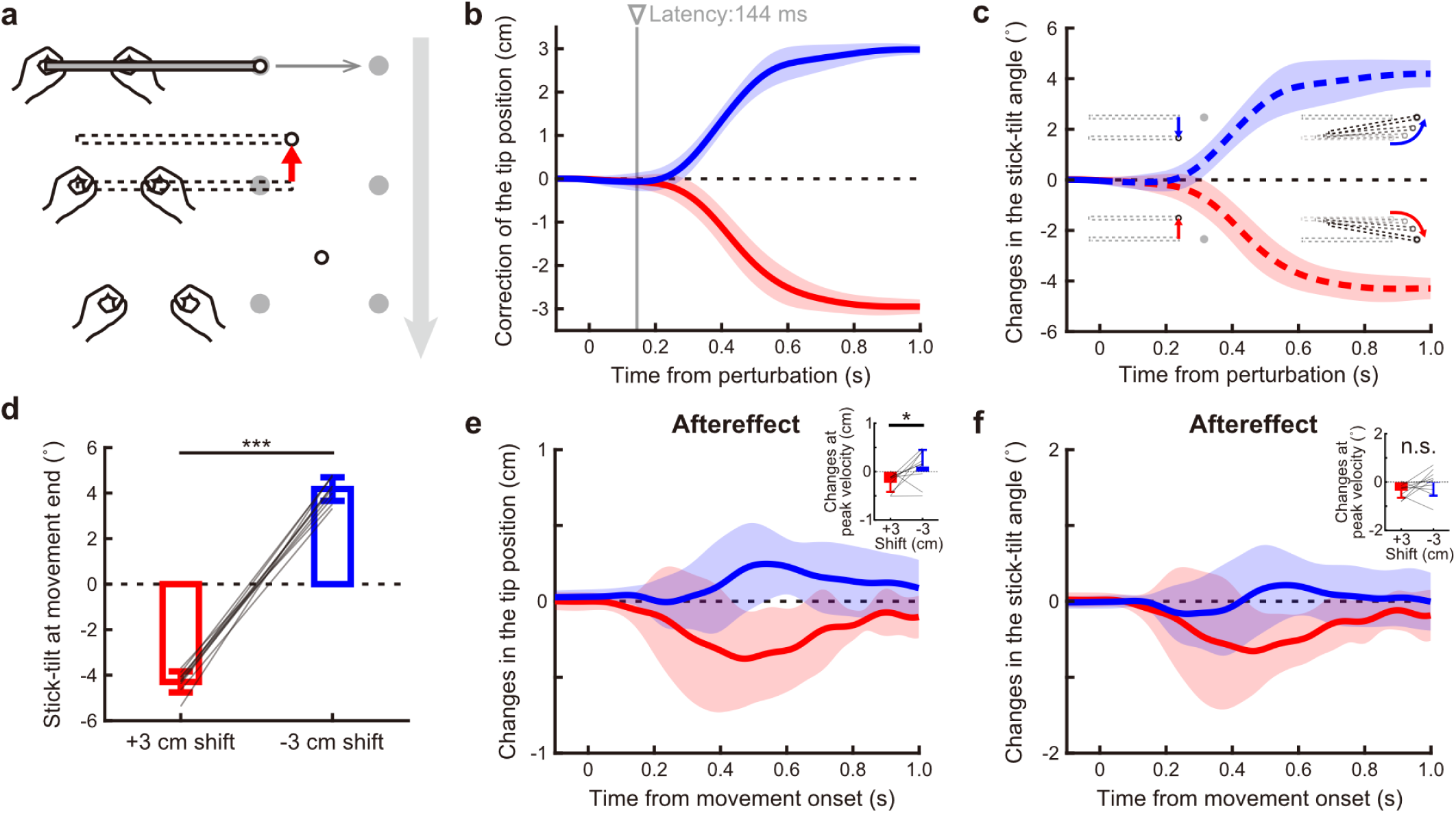
Corrective rapid response to the end-effector relevant perturbation in which the stick disappeared except for the right tip. (**a**) The stick position was shifted by ±3 cm and disappeared except for the tip. (**b**) Changes in the correction amount of the tip position error. The red (blue) plot shows average data when the stick was shifted by 3 cm (−3 cm). The participants rapidly corrected errors with a latency of 144 ms. Latency was derived from the time derivative of the position profiles (i.e., the velocity profiles). The shaded areas indicate the SD. (**c**) Changes in the hands-tilt angle show that the correction of the tip position was accompanied by a hands rotation. Insets depict the stick on the monitor, exaggeratedly drawn for clarity. (**d**) Changes in the hands-tilt angle at the movement end show that the hands rotation significantly accompanied the correction of the tip position (paired t-test, t(9) = −34.170, p < 0.001). The error bars indicate the SD. (**e**, **f**) In trials following the perturbation trials, biases of the tip position and stick-tilt angle were observed as aftereffects of the perturbations (insets: paired t-test, tip position: t(9) = −2.701, p = 0.024, stick-tilt angle: t(9) = −1.604, p = 0.143). *: p < 0.05, ***: p < 0.001.

### Rapid corrective responses to the end-effector irrelevant perturbation

To examine how the motor system responds to errors that do not directly affect task success, we introduced perturbations in an end-effector irrelevant dimension. Twenty-two participants (right-tip control: N = 11; left-tip control: N = 11) performed a perturbation phase consisting of 160 trials, including trials in which the stick angle around the controlled tip was unexpectedly rotated by ±6° during movement execution (Fig. 4a; Fig. S4a). Perturbations were applied pseudo-randomly once every four trials. Importantly, these perturbations altered the stick orientation without directly affecting the tip position. According to the minimal intervention principle, such end-effector irrelevant perturbations might be ignored, as they do not affect achievement of the task goal (Fig. 4b). Alternatively, if corrective responses are constrained by intrinsic coordination between end-effector relevant and end-effector irrelevant dimensions, correcting the stick-tilt error may systematically induce changes in tip position (Fig. 4c).

**Figure 4.**
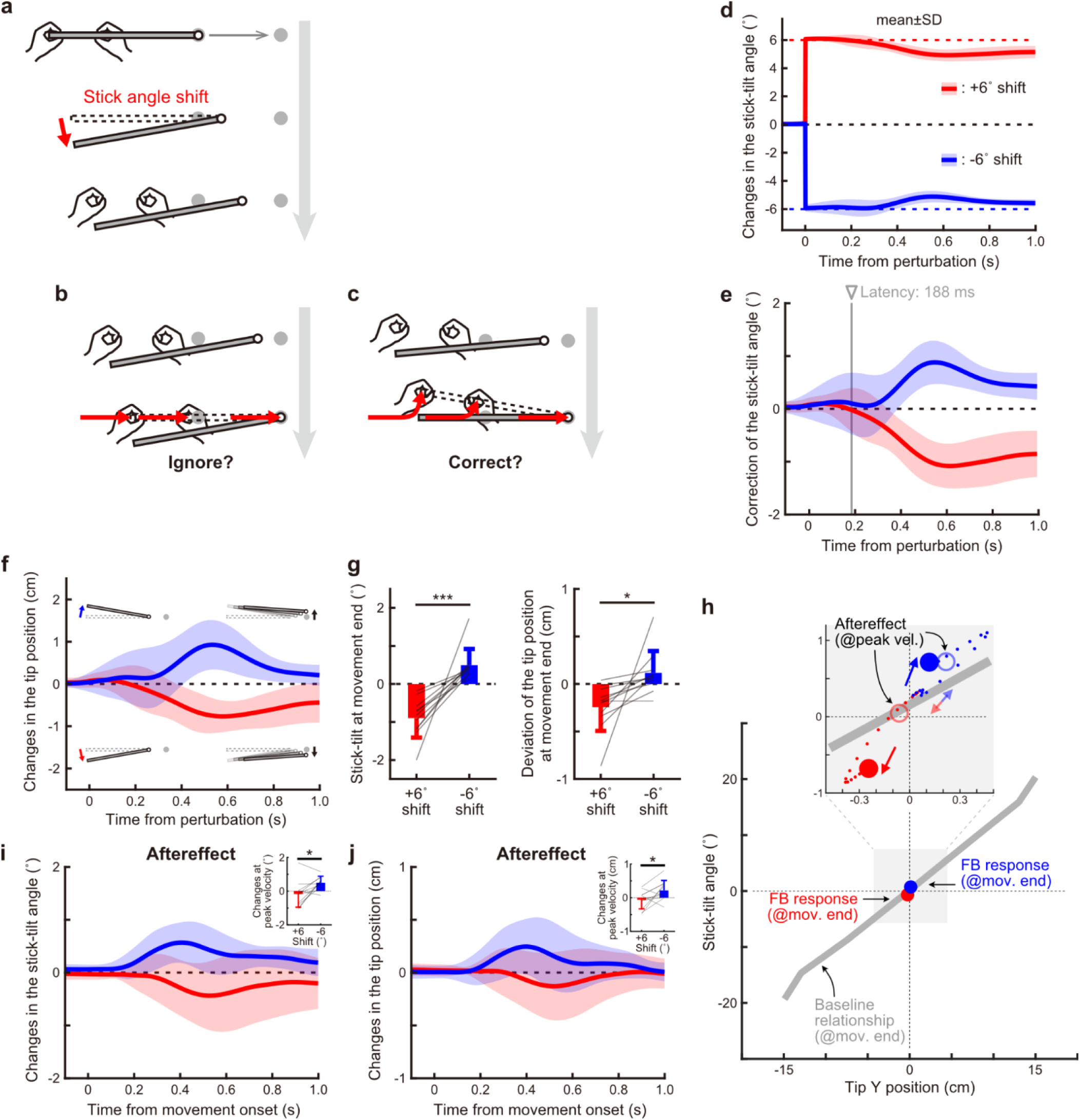
Corrective rapid response to the end-effector irrelevant perturbation. (**a**) In the perturbation trial of a stick angle shift, the stick suddenly rotated around the tip by +6 or −6° when the tip was 1 cm away from the starting point. The picture was exaggeratedly drawn for clarity. (**b**) Participants may not change the movement pattern because the perturbation did not affect the tip position. (**c**) Alternatively, they might correct for the stick-tilt angle while moving the tip. (**d**) The time-course changes in the stick-tilt angle. The red (blue) plot shows average data when the stick was rotated by 6° (−6°). The shaded areas indicate the SD. (**e**) Changes in the correction amount of the stick-tilt error. The participants rapidly corrected errors with a latency of 188 ms. Latency was derived from the time derivative of the angle profiles. (**f**) Changes in the tip position orthogonal to the target direction show that the correction of the stick-tilt angle was accompanied by a deviation of the tip during the reaching movement. Insets depict the stick on the monitor, exaggeratedly drawn for clarity. (**g**) Changes in the stick-tilt angle (*left*) and the deviation of the tip (*right*) at the movement end show that the tip position error significantly accompanied the compensative stick rotation (paired t-test, stick-tilt angle: t(10) =-5.333, p < 0.001; tip deviation: t(10) = −2.699, p = 0.022). The error bars indicate the SD. (**h**) A plane representing the relationship between the tip position and the stick-tilt angle. The formats are the same as Fig. 2h. The plots of the aftereffects are shown only in the inset. (**i, j**) In trials following the perturbation trials, biases of the stick-tilt angle and tip position were observed as aftereffects of the perturbations (insets: paired t-test, stick-tilt angle: t(10) = −2.999, p = 0.013, tip position: t(10) = −2.560, p = 0.028). *: p < 0.05, ***: p < 0.001.

Participants initiated corrective responses to the stick-tilt perturbation with a latency of approximately 188 ms (which was derived from the time derivatives of the angle profiles) (Figs. 4d, 4e), indicating rapid feedback control. Although the perturbation did not affect the tip position, corrective responses to the stick-tilt error were accompanied by systematic deviations of the tip trajectory (Fig. 4f). Specifically, counterclockwise corrections of the stick angle were associated with upward deviations of the tip, whereas clockwise corrections were associated with downward tip deviations. These coordination patterns closely matched those observed during baseline movements, indicating that corrective responses followed the same intrinsic relationship between stick tilt and tip position. At the end of the movement, both the corrected stick-tilt angle and the induced tip deviation remained significantly different from baseline (paired t-tests; stick-tilt angle: t(10) = −5.333, p < 0.001; tip deviation: t(10) = −2.699, p = 0.022; Fig. 4g). Similar results were observed when participants controlled the left tip of the stick (Figs. S4b–d). Plotting these movement patterns on the plane representing the relationship between the tip position and the stick-tilt angle revealed that they were still biased toward the baseline relationship (Fig. 4h).

In trials after the perturbation, participants exhibited aftereffects in stick angle, consistent with predictive compensation for the preceding perturbation (Fig. 4i; paired t-test, t(10) = −2.999, p = 0.013). Additionally, the aftereffect in tip position did not act to counteract the tip deviation induced by the feedback correction. Instead, tip position was biased in the same direction as the deviation that accompanied the corrective response in the previous trial (Fig. 4j; paired t-test, tip position: t(10) = −2.560, p = 0.028). This pattern was consistently observed regardless of whether participants controlled the right or left tip of the stick (Figs. S4b–d), indicating that it was independent of effector laterality. Again, the biases in movement patterns observed as aftereffects were consistent with the baseline relationship (Fig. 4h). Thus, even when the perturbation acted solely on an end-effector irrelevant dimension, the resulting learning did not compensate for cross-dimensional deviations generated during feedback correction.

Together, these results demonstrate that even errors that are irrelevant to task success evoke rapid and systematic corrective responses. Importantly, these responses propagate into end-effector relevant dimensions according to the same coordination constraints observed during unperturbed movements, indicating that fast feedback control in redundant motor systems is governed by intrinsic movement coordination structure rather than by selective correction of end-effector relevant errors alone.

### Dissociation between the stick and hand position alone does not elicit corrective responses

One possible alternative explanation for the corrective responses to stick-angle perturbations observed in the bimanual task is that they were driven not by errors in stick orientation per se, but by a positional discrepancy between the hand and the visually displayed stick. To test this possibility, we conducted control experiments using single-hand stick manipulation tasks. Twenty-four naïve participants performed either a left-hand task or a right-hand task (N = 12 each), following the same protocol as the bimanual task (baseline phase: 280 trials; perturbation phase: 160 trials). In the left-hand task, participants grasped the left tip of the virtual stick with their left hand (Fig. 5a), whereas in the right-hand task, participants grasped a position 15 cm from the left tip with their right hand (Fig. 5b). In both tasks, participants were instructed to move the right tip of the stick to a visual target. Crucially, in the single-hand tasks, participants could not control the stick orientation: the stick angle remained fixed at zero except during perturbation trials. During the perturbation phase, we applied the same stick-angle shifts (±6°) as in the bimanual task, which introduced a dissociation between the visual stick orientation and the actual hand position without affecting the tip position.

**Figure 5.**
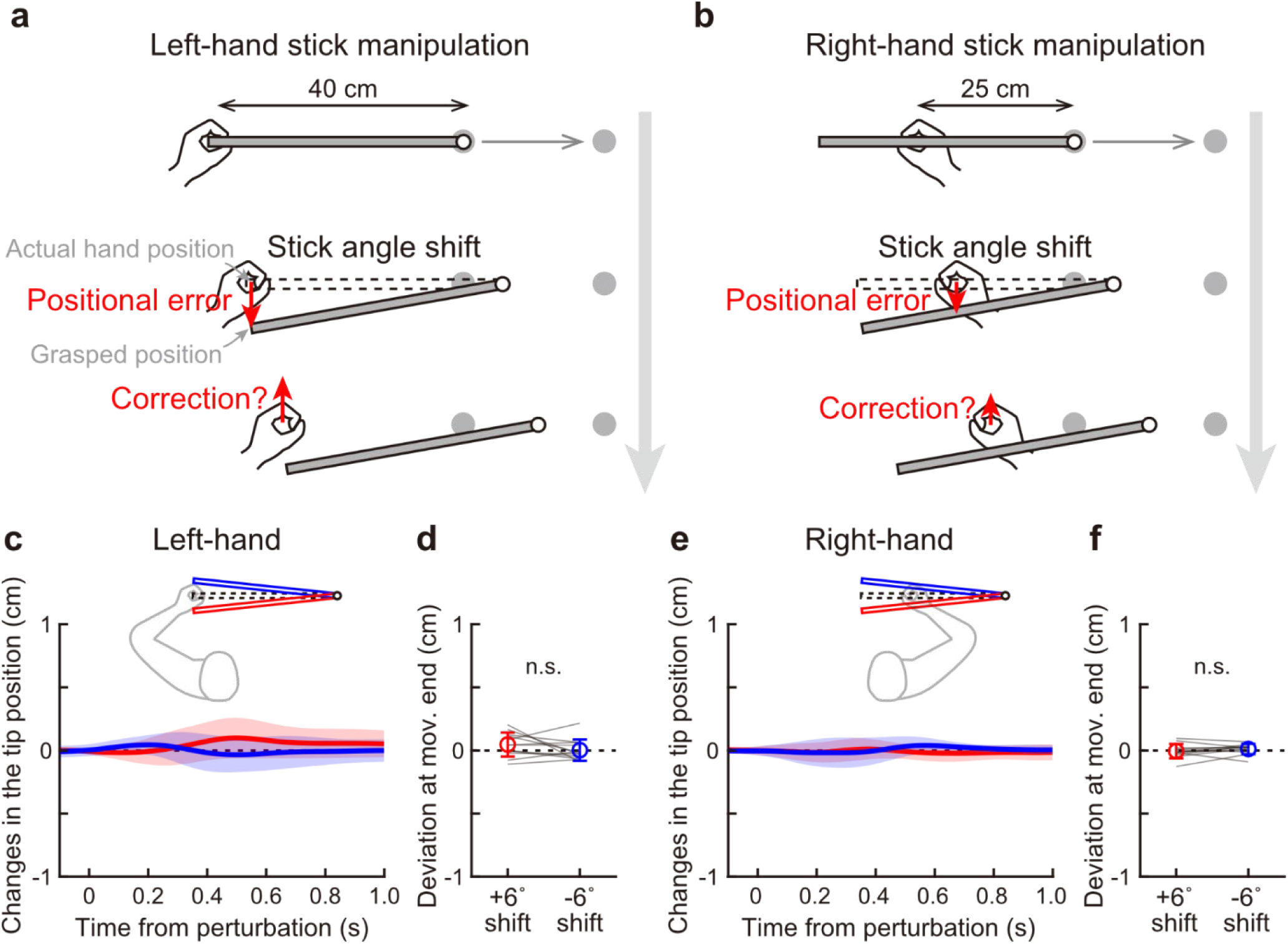
Single-hand stick manipulation task. (**a**, **b**) A total of 24 right-handers participated in the single-hand stick manipulation tasks. Twelve participants performed the left-hand task in which the tip was constantly displayed on 40 cm left of the left hand. The rest twelve participants performed the right-hand task in which the tip was constantly displayed on the 25 cm left of the right hand. During the perturbation phase following the baseline phase, the stick angle shift was introduced. Because the counterclockwise (clockwise) perturbation generates a backward (forward) error between the actual hand position and the grasped stick position, it may induce a corrective response by moving a hand forward (backward). (**c**) Changes in the tip position orthogonal to the target direction did not show corrective responses depending on the perturbation direction (left-hand condition). The red (blue) plot shows average data when the stick was rotated by 6° (−6°). (**d**) There was no significant difference in the tip position at the movement end (paired t-test, left-hand task, t(11) = 1.318, p = 0.214). (**e**, **f**) Similarly, the participants in the right-hand task showed no significant corrective response (paired t-test, right-hand task, t(11) = −1.103, p = 0.294).

If the positional discrepancy between the hand and the stick were sufficient to induce corrective responses, participants should have exhibited compensatory changes in movement even in the single-hand condition (Figs. 5a, 5b). However, no such corrective responses were observed: The time course of tip position showed no systematic deviation following the perturbation (Figs. 5c, 5e), and there was no significant difference in tip position at movement end for either perturbation direction (paired t-tests; left-hand task: t(11) = 1.318, p = 0.214, Fig. 5d; right-hand task: t(11) = −1.103, p = 0.294, Fig. 5f). These results indicate that a dissociation between the hand position and the visual representation of the stick alone is insufficient to elicit corrective responses. Instead, the corrective responses observed in the bimanual task likely depend on the ability to actively control and predict stick orientation. This supports the interpretation that corrective responses to stick-angle perturbations arise from intrinsic coordination between end-effector relevant and end-effector irrelevant dimensions, rather than from positional discrepancies at the level of the hand.

### End-effector irrelevant perturbations modulate corrective responses to end-effector errors

To further examine whether perturbations in end-effector–irrelevant dimensions influence the correction of end-effector errors, we simultaneously applied a stick position perturbation and a stick angle perturbation. In these trials, the displaced stick was additionally rotated either inward (inner shift) or outward (outer shift) relative to the direction of the tip displacement (Fig. 6a). According to the minimal intervention principle, corrective responses should primarily target the tip position error while ignoring the stick-angle perturbation, as the latter does not directly affect task success. Under this account, corrective responses should be similar across conditions. In contrast, if intrinsic coordination constraints couple end-effector relevant and end-effector irrelevant dimensions, the structure of the corrective response should depend on the relationship between the two perturbations. Specifically, in the inner-shift condition, the stick-angle correction biases the tip movement toward the target, potentially facilitating faster correction of the tip position. Conversely, in the outer-shift condition, the stick-angle correction biases the tip movement away from the target, potentially slowing the corrective response.

**Figure 6.**
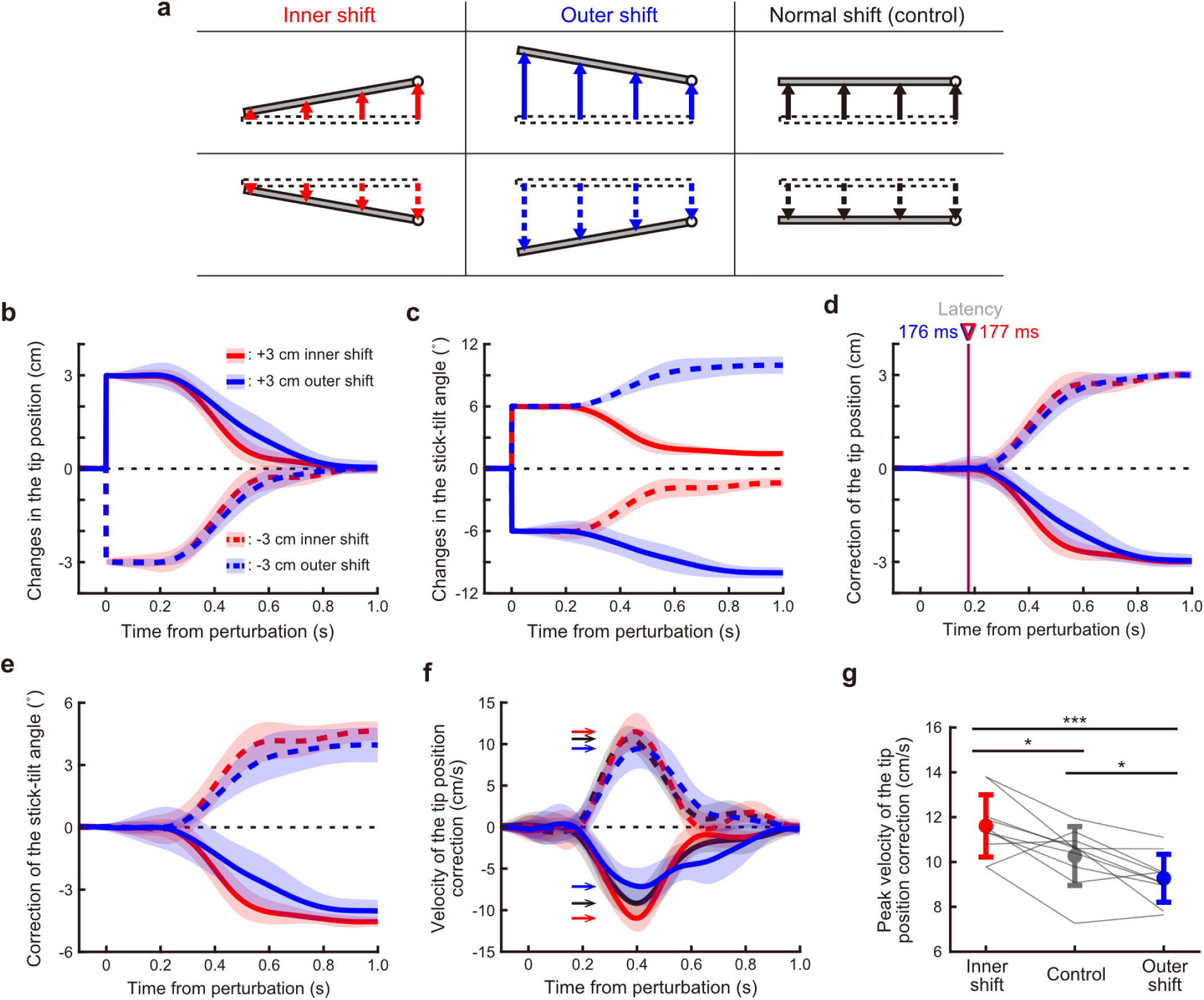
Effects of the end-effector irrelevant perturbation on the correction of the end-effector relevant error. (**a**) In the stick position shift conditions, we sometimes imposed the stick angle shift at the same as the stick position shift. Left: the combination of ±3 cm stick position shift and ±6° stick angle shift (inner shift). Middle: the combination of ±3 cm stick position shift and ∓6° stick angle shift (outer shift). Right: normal shift same as Fig. 2a (control). (**b**) The time-course changes in the tip position orthogonal to the target direction. The solid (dashed) plot shows averaged data when the stick was shifted by 3 cm (−3 cm), and the red (blue) plot shows averaged data of the inner shift (outer shift). The shaded areas indicate the SD. (**c**) The time-course changes in the stick-tilt angle. (**d**) Changes in the correction amount of the tip position error. Regardless of the direction of the stick angle shift, the participants rapidly corrected the tip position error with a latency of 176 ms (inner shift) or 177 ms (outer shift). Latency was derived from the velocity profiles. (**e**) Changes in the correction amount of the stick-tilt angle. (**f**) The correction velocity of the tip position was drawn by the derivative of the plots in c. Arrows in the figure indicate a peak of each velocity profile. The red, blue, and black plots indicate the result when the inner shift, outer shift, and normal shift were imposed, respectively. (**g**) The peak velocity of the tip position correction showed significant differences depending on the direction of the stick angle shift (a repeated-measures one-way ANOVA followed by a Bonferroni-holm post-hoc test, F(2, 18) = 19.785, p < 0.001, partial η2 = 0.687; inner shift vs. outer shift, t(9) = 7.004, p < 0.001, Cohen’s d = 1.846; inner shift vs. control, t(9) = 3.051, p = 0.028, Cohen’s d = 1.060; control vs. outer shift, t(9) = 2.969, p = 0.028, Cohen’s d = 0.786). *: p < 0.05, ***: p < 0.001.

Participants rapidly corrected the perturbed tip position in both conditions and appeared to largely ignore the stick-angle perturbation at the level of task outcome (Figs. 6b, 6c). Consistent with this, the latency of the corrective response to the tip position error did not differ between the inner- and outer-shift conditions (Fig. 6d). Despite similar response latencies, the kinematic structure of the corrective movements differed between conditions (Figs. 6d, 6e). These differences were reflected in the velocity profile of the tip correction (Fig. 6f). The peak correction velocity was significantly modulated by the direction of the stick-angle perturbation (a repeated-measures one-way ANOVA followed by a Bonferroni-holm post-hoc test, F(2, 18) = 19.785, p < 0.001, partial η2 = 0.687; inner shift vs. outer shift, t(9) = 7.004, p < 0.001, Cohen’s d = 1.846; inner shift vs. control, t(9) = 3.051, p = 0.028, Cohen’s d = 1.060; control vs. outer shift, t(9) = 2.969, p = 0.028, Cohen’s d = 0.786; Fig. 6g). Peak correction velocity was higher in the inner-shift condition than in both the control and outer-shift conditions, whereas correction was slower in the outer-shift condition. Similar effects were observed when participants controlled the left tip of the stick (Figs. S5a–d), indicating that this modulation was robust across task configurations.

These findings suggest that end-effector irrelevant perturbations influence end-effector correction not by changing the timing of corrective responses, but by shaping how corrective commands propagate through redundant motor degrees of freedom. Crucially, this pattern is consistent with the presence of intrinsic coordination constraints linking end-effector relevant and end-effector irrelevant dimensions.

## Discussion

The present study addressed a fundamental challenge in motor control posed by kinematic redundancy: how low-dimensional sensory errors are translated into coordinated changes across high-dimensional motor degrees of freedom. Traditional arm-reaching paradigms, in which end-effector position is uniquely determined by limb posture, do not help with this redundancy problem. By contrast, the redundant bimanual task used here allowed us to dissociate end-effector relevant and end-effector irrelevant dimensions and to reveal how intrinsic coordination constrains both feedback correction and trial-by-trial motor learning. We showed that visual errors propagate through the redundant motor system in a stereotyped manner, giving rise to fast corrective responses shaped by intrinsic coordination constraints linking end-effector relevant and end-effector irrelevant dimensions of movement. Crucially, this propagation was observed not only when perturbations directly affected task success, but also when perturbations acted solely on end-effector irrelevant dimensions. Moreover, trial-by-trial learning did not act to compensate for secondary deviations introduced by feedback correction across dimensions. Instead, aftereffects consistently reflected the structure of the corrective responses expressed in the preceding trial. Together, our results suggest that rapid motor corrections in redundant systems rely on stable coordination structures that enable efficient and reliable control in the face of unexpected perturbations, rather than on flexible re-optimization of motor commands at the moment an error occurs.

### Kinematic symmetry for the laterality of stick-tip manipulation

In the present stick-manipulation task, participants could in principle achieve the task either by translating both hands in parallel and/or by tilting the stick. Despite this redundancy, they consistently adopted a stereotyped strategy involving stick tilt from the beginning of the task. In the baseline phase, participants rotated the stick counterclockwise (clockwise) when aiming toward forward (backward) targets, a relationship previously shown to reflect minimization of the combined movement distance of both hands (Kobayashi & Nozaki, 2024).

This strategy leads to shorter movement distances for the hand farther from the controlled tip, consistent with an efficient allocation of movement across the two hands. This pattern could be interpreted as reflecting dominant-hand superiority, given that all participants were right-handed and dominant-arm control is often associated with superior movement accuracy (Sainburg, 2005). However, our results revealed a striking symmetry: when participants controlled the left tip of the stick, they exhibited the same stereotyped coordination pattern, with reduced movement of the right hand, despite its dominance. Thus, the baseline coordination structure was not anchored to handedness per se, but instead depended on which end of the stick served as the controlled end-effector.

Such symmetry is notable given extensive evidence for lateralized roles of the two arms in feedback responses and motor learning (Sainburg, 2005; Yokoi et al., 2014; Schaffer & Sainburg, 2021). Our findings instead align with prior work showing that, in redundant bimanual tasks, the motor system can flexibly exchange functional roles between the two hands depending on task context (Johansson et al., 2006). We therefore suggest that, in stick manipulation, the motor system dynamically assigns control roles to each hand based on the laterality of the controlled end-effector rather than on fixed hand dominance, enabling symmetric coordination under redundancy.

### Efficient rapid corrections to end-effector relevant perturbations in a redundant motor system

When the stick was suddenly shifted, participants rapidly corrected the resulting tip error by tilting the stick. The short corrective latency of approximately 160 ms indicates that this correction pattern was not the result of an explicit strategy, but rather an automatic feedback process (Scott, 2016). Notably, the corrective response involved a coordinated change across redundant degrees of freedom, resembling the stereotyped movement patterns observed during baseline movements. Tilting the stick to correct the tip position reduces the overall displacement of both hands compared to a strategy in which the hands move in parallel, suggesting that rapid feedback corrections exploit coordination patterns that are already efficient in the unperturbed task. The motor system might be able to rapidly reconfigure redundant effectors in an efficient manner even when confronted with unexpected perturbations.

Our previous work proposed that baseline coordination patterns serve as a scaffold that implicitly guides how corrective commands are distributed across redundant degrees of freedom during motor adaptation (Kobayashi & Nozaki, 2024). The present results indicate that this principle also extends to rapid feedback correction. Specifically, when the tip was displaced backward (forward), participants corrected it forward (backward) while rotating the stick counterclockwise (clockwise), preserving the baseline relationship between tip movement direction and stick-tilt angle.

This consistency between baseline coordination and rapid feedback responses parallels previous findings that both motor adaptation and feedback control are shaped by common coordination structures (Hayashi et al., 2016). Importantly, this guidance by baseline coordination differs fundamentally from trial-by-trial optimization of movement efficiency. Instead, it suggests that rapid corrections are generated by recruiting pre-existing coordination structures that constrain how corrective commands propagate through the redundant motor system.

### Why end-effector irrelevant perturbations cannot be ignored in redundant motor control

We introduced an end-effector irrelevant perturbation by rotating the stick around the controlled tip, which did not directly affect the end-effector position. In principle, such perturbations could be ignored without compromising task performance. According to the minimal intervention principle, the motor system should therefore be indifferent to errors confined to task-irrelevant dimensions. Contrary to this prediction, participants reflexively corrected the stick-tilt error during movement, indicating that the motor system responded to perturbations even when they did not affect task success.

One possible explanation is that corrective responses were constrained by intrinsic coordination linking end-effector relevant and end-effector irrelevant dimensions of movement (Kobayashi & Nozaki, 2024). In the bimanual task, baseline movements exhibited a tight coupling between tip movement direction and stick-tilt amplitude. Moreover, corrections to end-effector relevant perturbations consistently followed this baseline relationship. We propose that this coordination constraint also governs responses to perturbations confined to end-effector irrelevant dimensions. Even when task success is unaffected, deviations from the baseline coordination structure may be detected and corrected, leading to systematic changes in both stick orientation and tip position. This framework explains why end-effector irrelevant perturbations could not be selectively ignored in the redundant bimanual task, while no such responses emerged in the single-hand condition, where no intrinsic coupling between stick orientation and tip movement existed.

Another possibility is that the motor system predicts the state of the entire manipulated object, rather than focusing exclusively on the end-effector. This idea is consistent with the notion that humans tend to experience a sense of agency over objects whose motion is caused by their own actions (Haggard, 2017). Perturbations to stick orientation may have induced sensory prediction errors between proprioceptive signals from the hands and visual information about the stick, prompting corrective responses. Such sensory prediction errors are known to drive both visuomotor adaptation (Wolpert et al., 1995; Flanagan & Rao, 1995; Mazzoni & Krakauer, 2006; Schaefer et al., 2012; Kim et al., 2019) and feedback corrective response (Nashed et al., 2012; Franklin et al., 2016). However, this explanation was not supported by our control experiments: In the single-hand stick manipulation tasks, in which a comparable dissociation between visual stick orientation and hand position was introduced, participants showed no corrective responses. This result argues against sensory prediction error between proprioceptive and visual signals as the primary driver of the observed responses.

### Visuomotor mapping-dependent error correction

The single-hand stick manipulation task showed that a positional discrepancy between the hand and the stick alone is insufficient to elicit corrective responses. Because the stick remained horizontal regardless of the direction of hand movement in this task, participants lacked a visuomotor mapping linking their movements to stick rotation. It is generally believed that the motor system maintains an internal relationship between motor commands and their outcomes (Schmidt, 1975) and that the same control schema can be recruited as a visuomotor mapping for error correction (Hayashi et al., 2016). Our findings suggest that the converse does not necessarily hold: the presence of a visual error alone does not trigger correction unless an appropriate control schema has been formed. This may highlight the importance of experiencing a variety of trial-and-error patterns when learning a new motor skill.

### Interaction between end-effector relevant and end-effector irrelevant error corrections

To examine how end-effector relevant and end-effector irrelevant errors interact during feedback correction, we simultaneously perturbed the tip position and stick orientation. Participants rapidly corrected the tip position error, consistent with prioritization of end-effector errors. However, corrective responses to stick-tilt angle occurred concurrently, and the resulting movement patterns followed the baseline coordination structure linking tip movement direction and stick-tilt angle. Although the latency of tip position correction was not affected by the presence of stick-tilt perturbation, the velocity profile of the correction was significantly modulated depending on whether the stick rotation facilitated or opposed the tip correction. These results indicate that, while end-effector errors are prioritized in terms of when corrective responses are initiated, intrinsic coordination constraints shape how corrective commands are distributed across redundant degrees of freedom during feedback control.

In summary, we show that the motor system exploits a baseline coordination structure to propagate low-dimensional visual errors into high-dimensional movement degrees of freedom, giving rise to rapid corrective responses. These corrections are not generated through flexible, moment-to-moment re-optimization, but are shaped by intrinsic coordination constraints that couple end-effector relevant tip motion with end-effector irrelevant stick tilt. In addition, trial-by-trial learning did not act to compensate for secondary deviations introduced by feedback correction across dimensions. Our findings indicate that both feedback correction and learning are governed by the same intrinsic coordination structure.

## Materials and Methods

### Participants

A total of 65 healthy adults (33 females and 32 males, aged 25.2 ± 7.2 years old; mean ± SD) participated in the study. The participants were recruited at The University of Tokyo or by the online recruiting system (https://www.jikken-baito.com). All participants were right-handed according to the laterality score (84.0 ± 16.1; mean ± SD) obtained by the Edinburgh Handedness Inventory (Oldfield, 1971). They gave written informed consent to an experimental protocol approved by the ethical committee of The University of Tokyo (#19-225). As described below, the participants were randomly assigned to either of four conditions (laterality of workspace; type of perturbation).

### Experimental apparatus

The participants performed reaching movements using a virtual stick displayed on a horizontal screen with two manipulandum handles (KINARM End-Point Lab; Kinarm, Kingston, Canada) (Scott, 1999) (Fig. 1a). The participants could not directly see their arms and handles because the screen occluded them. Instead, the screen displayed a starting position, a target (1.4 cm in diameter), a virtual stick (length: 40 cm, width: 0.5 cm). A total of 21 participants held the left tip of the stick with their left hand (left handle) and a position 15 cm away from the left hand with their right hand (right handle), and they were asked to control a white circle (1 cm in diameter) on the right tip of the stick (Fig. 1a). The remaining 20 participants performed a similar task in the reversed workspace (see Fig. S1a). The distance between the handles was fixed using a strong elastic force produced by the manipulanda (spring constant: 2000 N/m). The positions and velocities of the handles were analogue/digitally converted at 1.129 kHz and then recorded at 1 kHz for offline analysis.

### Baseline phase

All experiments comprised a baseline phase (280 trials), which was common to all conditions, and a perturbation phase (stick position shift condition: 360 trials, or stick angle shift condition: 160 trials). The baseline phase was performed to investigate the basic strategy of how the participants moved the stick to reach the targets with the right (or left) tip. At the beginning of each trial, the participants were asked to move the stick to its home position (hands and stick in Fig. 1c). After maintaining the home position for 0.5−1 s, a gray target was pseudo-randomly chosen from seven possible targets (0°: horizontal direction, ±30°, ±60°, ±90°; Fig. 1c) appearing 15 cm away from the initial right (or left) tip position of the stick. After waiting for another 0.5−1 s, the color of the target changed to pink, indicating that they should immediately start moving the tip to the target. The participants were instructed to maintain the peak velocities of the tip as constant as possible across the trials. When the tip reached the target, a feedback message, “FAST” or “SLOW,” was presented on the monitor if the peak speed was higher or lower than the range of 400−550 mm/s, respectively. After maintaining the tip at the target for 0.5 s, the visual feedback of the stick and target disappeared. Then, the handles automatically returned to their home position, and the next trial began.

### Perturbation phase

During the subsequent perturbation phase, the target appeared only in the 0° direction, and visual perturbations were sometimes imposed as probe trials. The home positions, size of the visual feedback and the target, inter-trial interval, and criteria for the velocity feedback were the same as those in the baseline phase. The laterality of the workspace remained unchanged through the experiment.

A total of 19 participants (right workspace: N = 10, left workspace: N = 9) were assigned to stick position shift condition (360 trials). To investigate how the motor system changes movement patterns when encountering an end-effector relevant error, we introduced end-effector relevant perturbations visually shifting the stick position. The participants randomly experienced either of eight perturbations (type of shifts: 4 * shift direction: 2) every three trials (each perturbation: 15 trials). First, the entire stick was displaced perpendicularly to the target direction (Fig. 2a). Second, the stick was shifted and disappeared except for the tip (Fig. 3a). Third, the stick was shifted and rotated in the inner direction by 6° (Fig. 6a, left). Fourth, the stick was shifted and rotated in the outer direction by 6° (Fig. 6a, middle). Importantly, the amount by which the tip position was shifted (3 cm) and the timing at which the shift was imposed (when the tip was 1 cm away from the home position) were common to all perturbations. If the motor system was indifferent to the end-effector irrelevant information, it should have shown the same correction pattern to these four perturbations.

Furthermore, to directly test whether the motor system ignores the end-effector irrelevant error, we introduced end-effector irrelevant perturbations by rotating the stick around the tip (Fig. 4a). A total of 22 participants (right workspace: N = 11, left workspace: N = 11) were assigned to this stick angle shift condition (160 trials). The participants randomly experienced either of two perturbations (rotation direction: 2) every four trials (each perturbation: 20 trials). The stick suddenly rotated around the tip by 6° when the tip position was 1 cm away from the starting point. The participants did not need to respond to the stick rotation because this perturbation did not change the tip position.

### Single-hand task

To investigate whether dissociation of the stick and holding position induced the responses to the stick angle shift, we conducted single-hand stick manipulation tasks and introduced the stick angle shift. Newly recruited participants performed a left-hand task (N =12) or right-hand task (N = 12). The home positions, target configurations, size of the visual feedback and the target, inter-trial interval were the same as those in the bimanual task (baseline phase (280 trials) + perturbation phase (160 trials)). Participants in the left-hand task held the left tip of the virtual stick with a left hand (Fig. 5a), whereas participants in the right-hand task held a position 15 cm away from the left tip with a right hand (Fig. 5b). Because they could not control the angle of the stick-tilt, the stick on the monitor was constantly horizontal except in the perturbation trials.

### Data analysis

Data analyses were performed using MATLAB R2025a (Mathworks, Natick, Massachusetts). Before the analysis, the behavioral data were low-pass filtered (10 Hz, fourth-order Butterworth). No participant was rejected from the analysis.

The movement onset was detected as a time point when the tip velocity started to exceed 1% of the peak velocity at least for 0.1 s. Similarly, the movement offset was defined as a time when the tip velocity started to fall below the same criteria. When calculating feedback responses and their aftereffects, to correct for participant-specific movement biases, we subtracted the mean data of the pre-perturbation null trials from the perturbation and aftereffect trials for each participant.

Latency of the corrective responses to the visual perturbations was defined as the time point at which responses to opposite perturbations split, and was estimated using a receiver-operator characteristic (ROC) knee method (Pruszynski et al., 2008; Yang et al., 2011; Crevecoeur et al., 2012). To increase detection sensitivity, latency was derived from the time derivatives of the position or stick-tilt angle profiles. The area under the ROC curve (AUC) for each time step was generated to calculate the probability that two responses to the distinct perturbations could be distinguished. A discrimination time point was defined as the first time point when the AUCs were < 0.25 or > 0.75 for five consecutive samples. We then calculated the fitting line to 30 samples (30 ms) of the AUCs, centered on the discrimination time point. Latency was defined as a time point when the fitting line intersected with the chance level (AUC = 0.5) line.

### Statistical analysis

To statistically evaluate the behavioral data, we conducted *t*-tests and repeated-measures one-way ANOVAs using MATLAB and the software package JASP (http://jasp-stats.org). We used Mauchly’s test to check sphericity of the data when performing ANOVAs. If sphericity was violated, Greenhouse-Geisser correction was applied to correct *p* values.

With respect to the feedback response, we quantified the response at the movement endpoint, where the response to the perturbation had fully accumulated. In contrast, with respect to the learning response, we quantified the response at peak velocity rather than at the endpoint to avoid contamination by online corrective components during the movement.

## Acknowledgments

We thank members of the Nozaki laboratory for their helpful comments and suggestions, Yudai Suzuki for performing preliminary experiments, Asako Munakata and Mayumi Yoda for coordinating experiments. This study was supported by a grant from the Japan Society for the Promotion of Science Research Fellowships for Young Scientists to T.K. (20J13734, 25KJ0444) and a KAKENHI to D.N. (17H00874, 21H04860) and to T.K. (23K10739).

## Author Contributions

Conceptualization, Methodology, Investigation, Formal Analysis, Writing: TK, DN, Supervision: DN.

---

**Figure S1.**
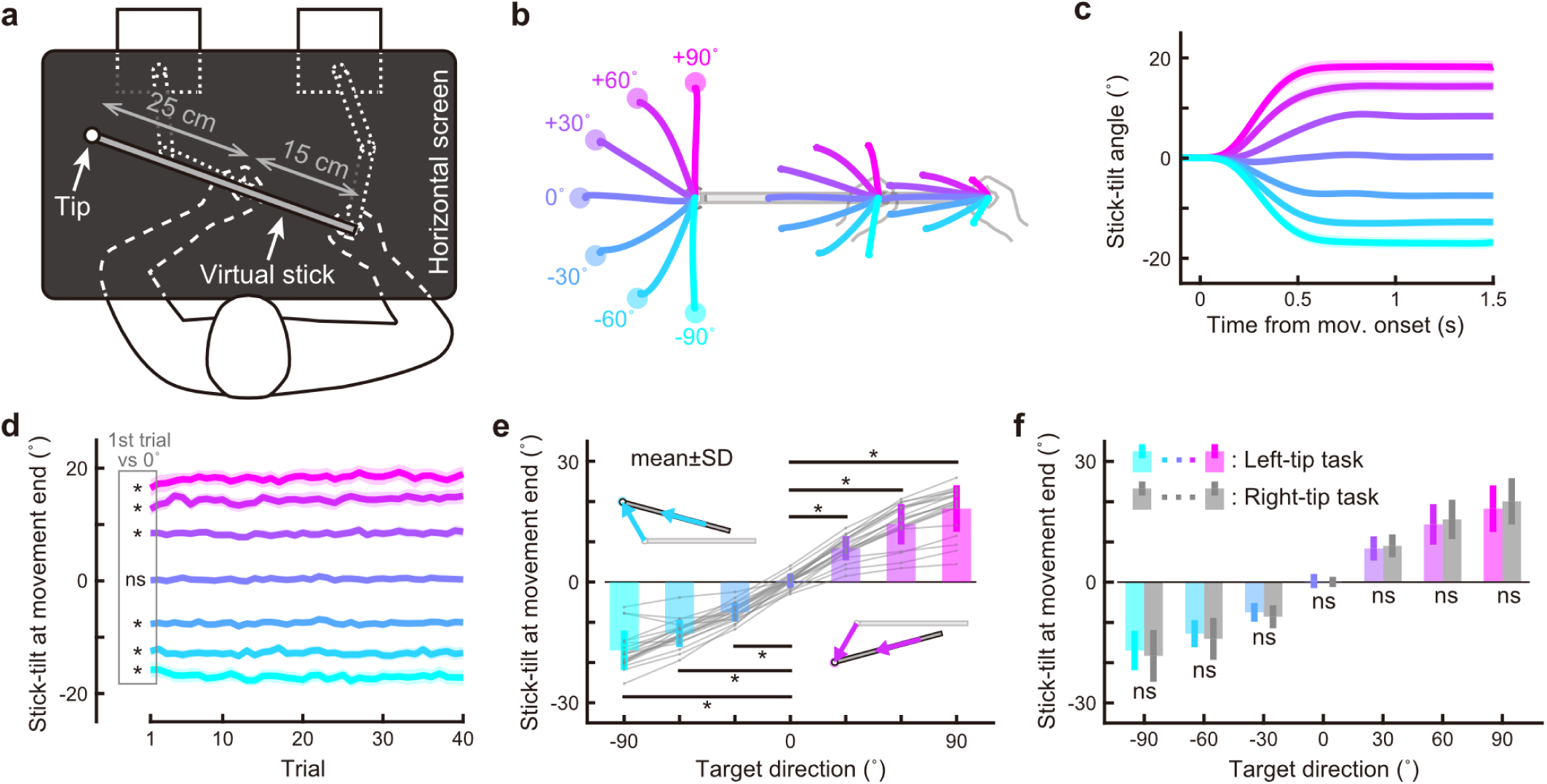
Kinematics to move the left tip of the stick with both hands. (**a**) Participants held handles with both hands and moved the left tip of a virtual stick on the screen. (**b**) During the baseline phase, participants were instructed to move the tip from a home position (a pale picture of the stick) to a target that appeared in one of seven directions. Movement trajectories of the tip, left hand, and right hand were drawn over the diagram (averaged across participants). (**c**) The time-course of the stick-tilt angle during the reaching movements. They achieved the task while tilting the stick. Each color represents the direction of the target, as shown in **b**. The shaded areas indicate SEM. (**d**) Trial-to-trial changes in the stick-tilt angle at the movement end. They adopted the strategy of tilting the stick from the beginning of the experiment (t-test compared with zero, |t(19)| > 10.9, Bonferroni corrected p < 0.001). The shaded areas indicate SEM. (**e**) The stick-tilt angle at the movement end significantly differs compared to that when aiming at a horizontal target (paired *t*-test, |t(19)| > 13.3, Bonferroni corrected p < 0.001). Gray plots indicate individual data. The error bars mean SD. (**f**) Comparisons between the right-tip control condition and left-tip control condition. There was no significant difference between the stick-tilt angles (t-test, |t(39)| < 1.32, Bonferroni corrected p > 0.19). *: p < 0.001.

**Figure S2.**
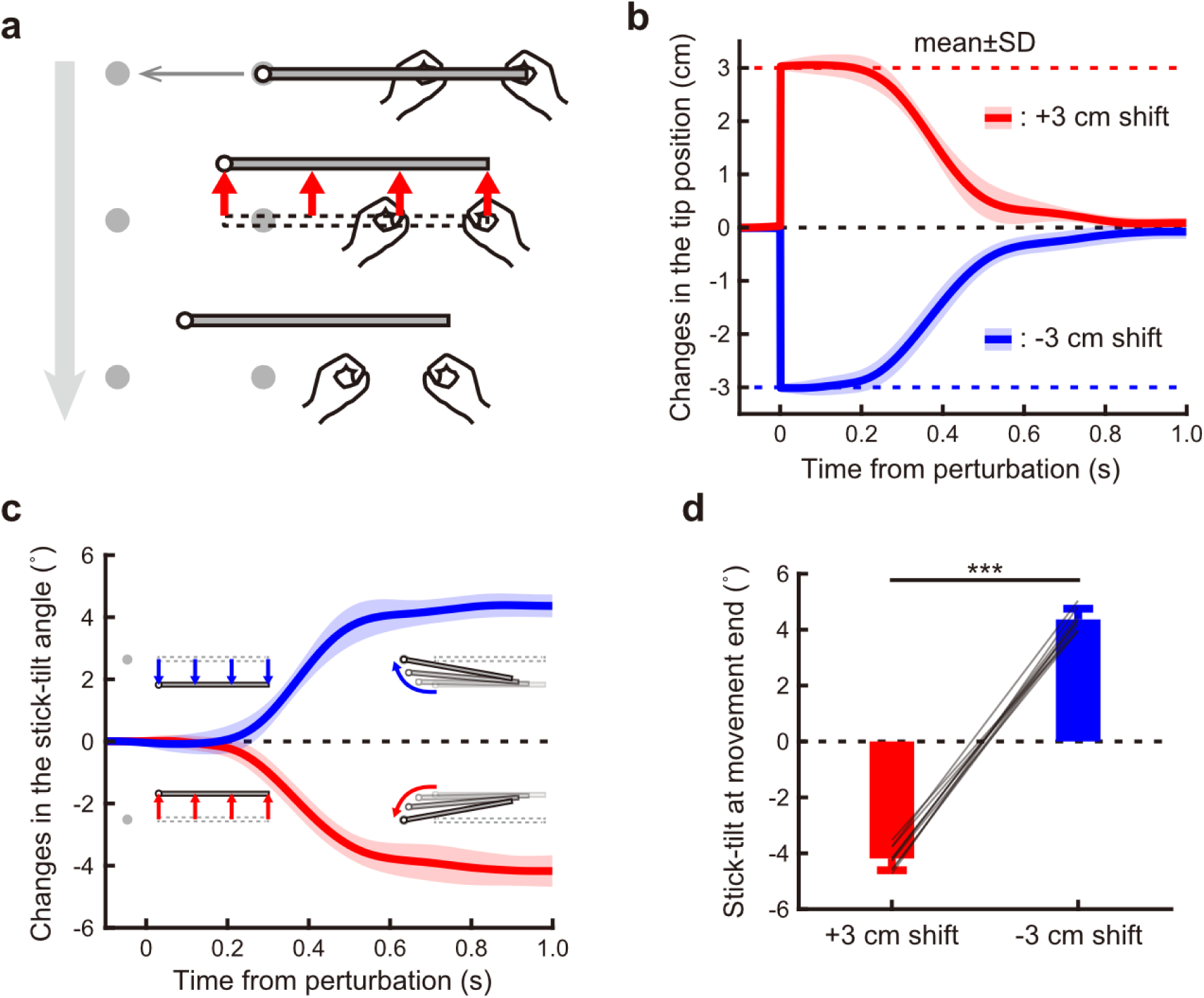
Corrective rapid response to the end-effector relevant perturbation in the left-tip control condition. (**a**) The stick position was shifted by ±3 cm. (**b**) The time-course changes in the tip position orthogonal to the target direction. The formats are the same as Fig. 2. (**c**) Changes in the stick-tilt angle show that the correction of the tip position was accompanied by a stick rotation. Insets depict the stick on the monitor, exaggeratedly drawn for clarity. (**d**) Changes in the stick-tilt angle at the movement end show that the stick rotation significantly accompanied the correction of the tip position (paired t-test, t(8) = −40.636, p < 0.001). The error bars indicate the SD. ***: p < 0.001.

**Figure S3.**
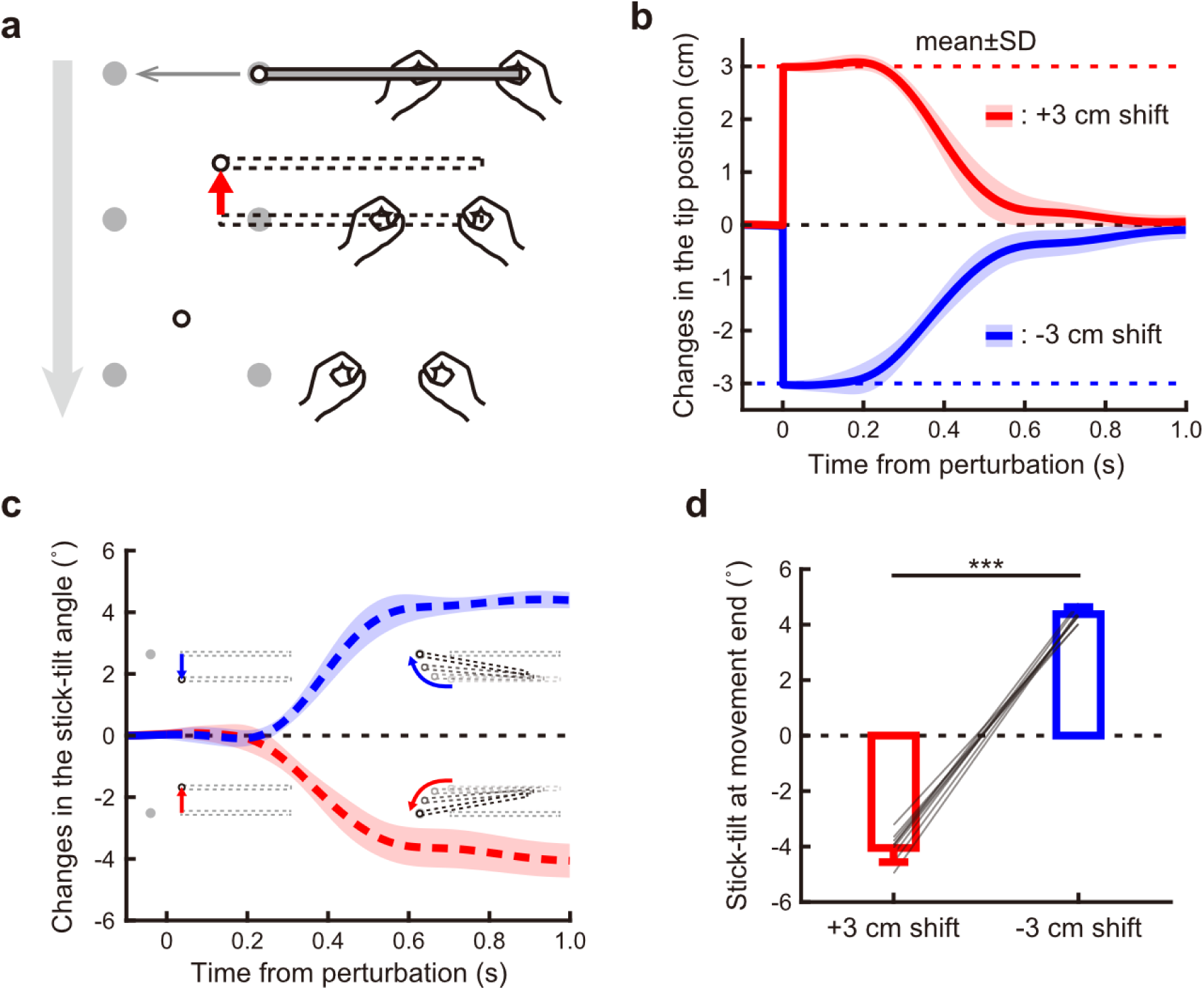
Corrective rapid response to the end-effector relevant perturbation in which the stick disappeared except for the left tip. (**a**) The stick position was shifted by ±3 cm and disappeared except for the tip. (**b**) The time-course changes in the tip position orthogonal to the target direction. The formats are the same as the Fig. 2. (**c**) Changes in the hands-tilt angle show that the correction of the tip position was accompanied by a hands rotation. Insets depict the stick on the monitor, exaggeratedly drawn for clarity. (**d**) Changes in the hands-tilt angle at the movement end show that the hands rotation significantly accompanied the correction of the tip position (paired t-test, t(8) = −36.632, p < 0.001). The error bars indicate the SD.

**Figure S4.**
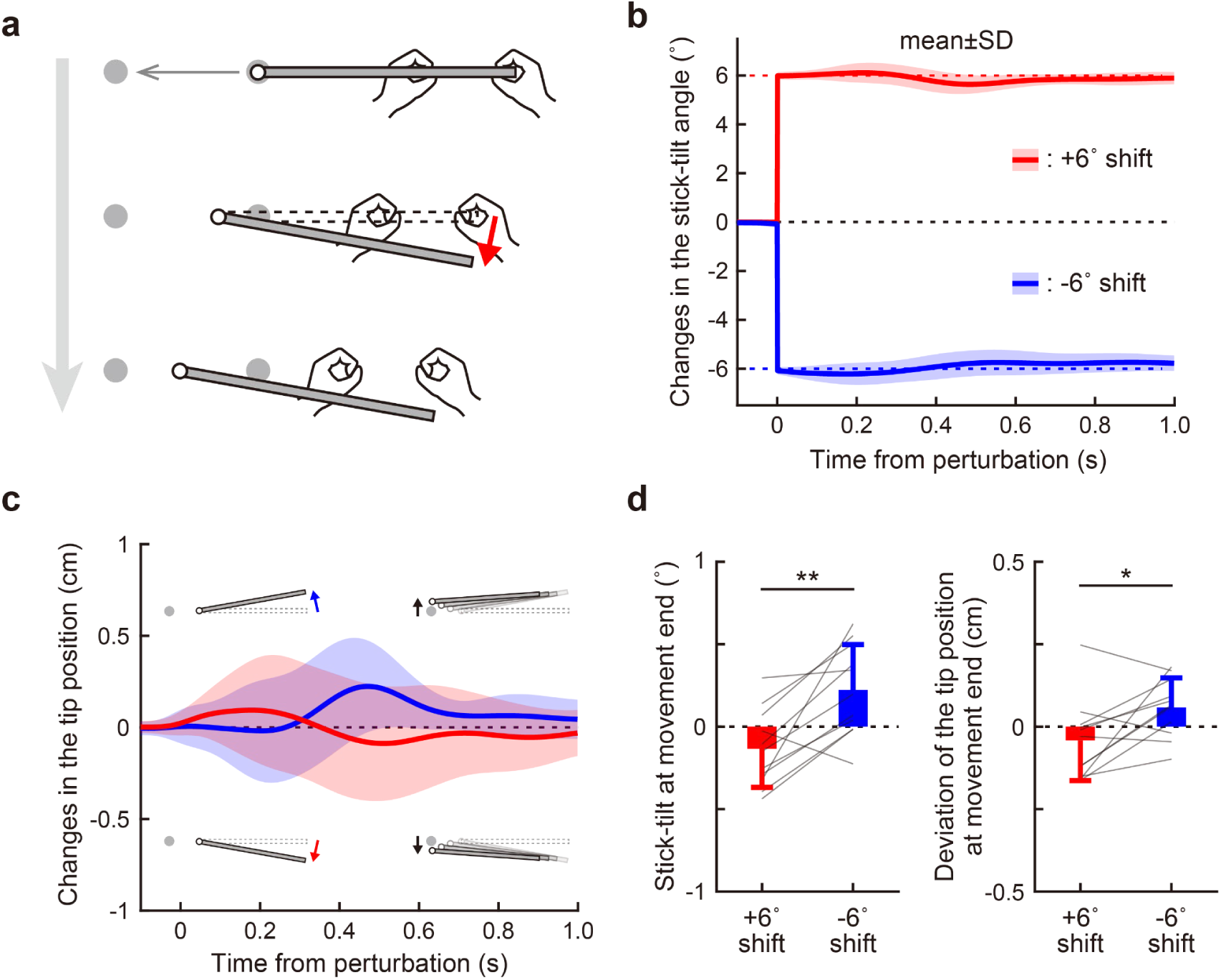
Corrective rapid response to the end-effector irrelevant perturbation in the left-tip control condition. (**a**) The stick suddenly rotated around the tip by 6° when the tip was 1 cm away from the starting point. The picture was exaggeratedly drawn for clarity. (**b**) In the perturbation trial of a stick angle shift, the stick was suddenly rotated around the tip by +6 or −6° when the tip was 1 cm away from the starting point. The picture was exaggeratedly drawn for clarity. The formats are the same as Fig. 4. (**c**) Changes in the tip position orthogonal to the target direction show that the correction of the stick-tilt angle was accompanied by a deviation of the tip during the reaching movement. Insets depict the stick on the monitor, exaggeratedly drawn for clarity. (**d**) Changes in the stick-tilt angle (*left*) and the deviation of the tip (*right*) at the movement end show that the tip position error significantly accompanied the compensative stick rotation (paired t-test, stick-tilt angle: t(10) =-4.195, p = 0.002; tip deviation: t(10) = −2.668, p = 0.024). *: p < 0.05, **: p < 0.01.

**Figure S5.**
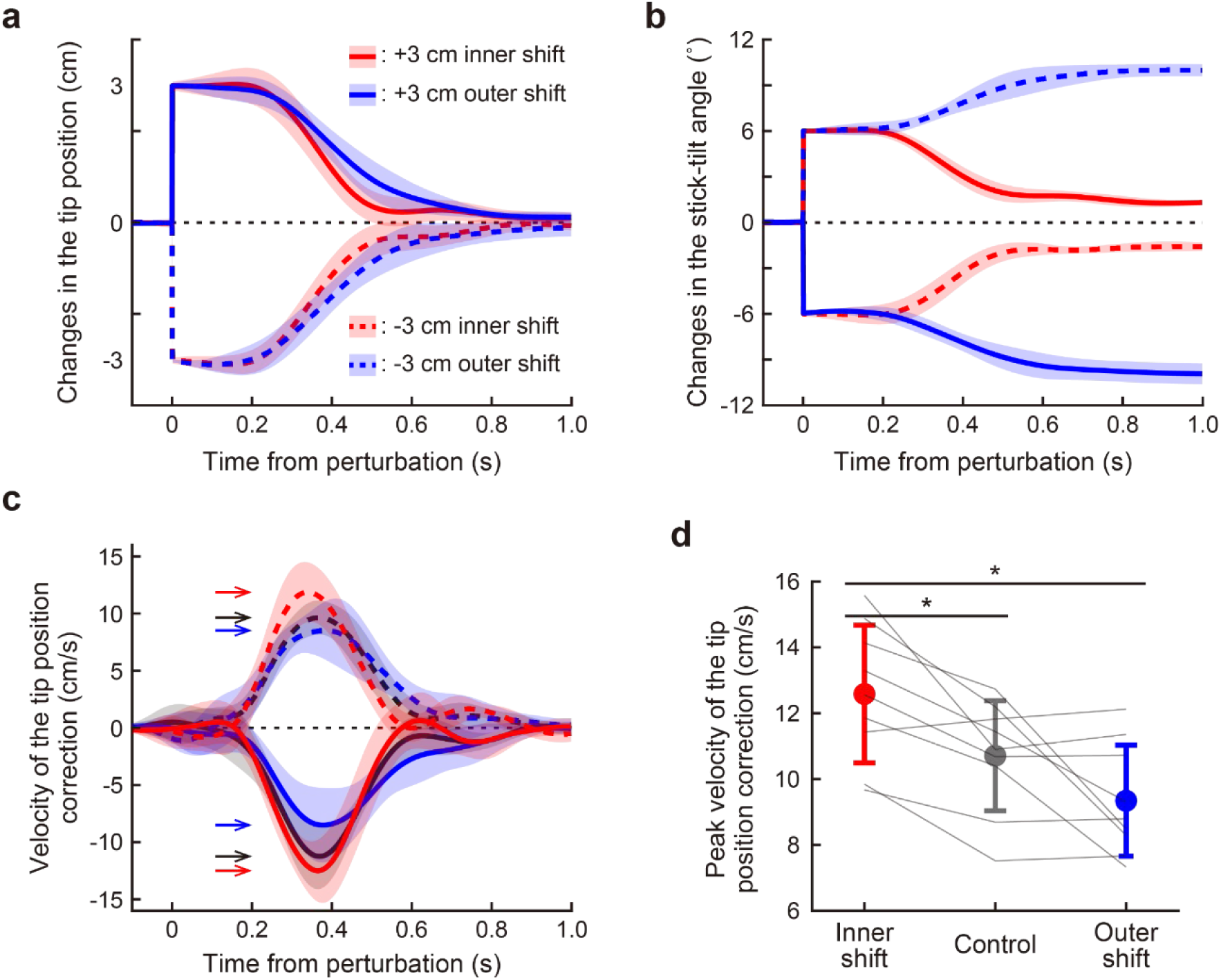
Effects of the end-effector irrelevant perturbation on the correction of the end-effector relevant error in the left-tip control condition. (**a**) The time-course changes in the tip position orthogonal to the target direction. The formats are the same as Fig. 6. (**b**) The time-course changes in the stick-tilt angle. (**c**) The correction velocity of the tip position. Arrows in the figure indicate a peak of each velocity profile. (**d**) The peak velocity of the tip position correction showed significant differences depending on the direction of the stick angle shift (a repeated-measures one-way ANOVA followed by a Bonferroni-holm post-hoc test, F(2, 16) = 12.686, p < 0.001, partial η^2^ = 0.613; inner shift vs. outer shift, t(8) = 4.130, p = 0.010, Cohen’s d = 1.774; inner shift vs. control, t(8) = 4.098, p = 0.010, Cohen’s d = 1.026; control vs. outer shift, t(8) = 2.090, p = 0.070, Cohen’s d = 0.748). *: p < 0.05.

## References

Morasso P. 1981. Spatial Control of Arm Movements. Experimental Brain Research 42:223−227.

Flash T, Hogan N. 1985. The Coordination of Arm Movements: An Experimentally Confirmed Mathematical Model. Journal of Neuroscience 5:1688−1703.

Gordon J, Ghilardi MF, Ghez C. 1994. Accuracy of planar reaching movements I. Independence of direction and extent variability. Experimental Brain Research 99:97–111.

Shadmehr R, Mussa-Ivaldi FA. 1994. Adaptive Representation of Dynamics During Learning of a Motor Task. Journal of Neuroscience 14:3208–3224.

Thoroughman KA, Shadmehr R. 1999. Electromyographic Correlates of Learning an Internal Model of Reaching Movements. Journal of Neuroscience 19:8573–8588.

Krakauer JW, Pine ZM, Ghilardi M-F, Ghez C. 2000. Learning of Visuomotor Transformations for Vectorial Planning of Reaching Trajectories. Journal of Neuroscience 20:8916–8924.

Scott SH. 2016. A Functional Taxonomy of Bottom-Up Sensory Feedback Processing for Motor Actions. Trends in Neurosciences 39:512−526.

Goodale MA, Pélisson D, Prablanc C. 1986. Large adjustments in visually guided reaching do not depend on vision of the hand or perception of target displacement. Nature 320:748−750.

Pélisson D, Prablanc C, Goodale MA, Jeannerod M. 1986. Visual control of reaching movements without vision of the limb - II. Evidence of fast unconscious processes correcting the trajectory of the hand to the final position of a double-step stimulus. Experimental Brain Research 62:303−311.

Desmurget M, Epstein CM, Turner RS, Prablanc C, Alexander GE, Grafton ST. 1999. Role of the posterior parietal cortex in updating reaching movements to a visual target. Nature Neuroscience 2:563−567.

Day BL, Lyon IN. 2000. Voluntary modification of automatic arm movements evoked by motion of a visual target. Experimental Brain Research 130:159−168.

Pisella L, Gréa H, Tilikete C, Vighetto A, Desmurget M, Rode G, Boisson D, Rossetti Y. 2000. An ‘automatic pilot’ for the hand in human posterior parietal cortex: Toward reinterpreting optic ataxia. Nature Neuroscience 3:729−736.

Dimitriou M, Wolpert DM, Franklin DW. 2013. The Temporal Evolution of Feedback Gains Rapidly Update to Task Demands. Journal of Neuroscience 33:10898−10909.

Franklin DW, Reichenbach A, Franklin S, Diedrichsen J. 2016. Temporal Evolution of Spatial Computations for Visuomotor Control. Journal of Neuroscience 36:2329−2341.

Wei K, Körding K. 2009. Relevance of Error: What Drives Motor Adaptation? Journal of Neurophysiology 101:655−664.

Kasuga S, Telgen S, Ushiba J, Nozaki D, Diedrichsen J. 2015. Learning feedback and feedforward control in a mirror-reversed visual environment. Journal of Neurophysiology 114:2187−2193.

Bernstein NA, Latash MA, Turvey MT. 1996. Dexterity and Its Development. Psychology Press.

Todorov E, Jordan MI. 2002. Optimal feedback control as a theory of motor coordination. Nature Neuroscience 5:1226−1235.

Todorov E. 2004. Optimality principles in sensorimotor control. Nature Neuroscience 7:907−915.

Scott SH. 2004. Optimal feedback control and the neural basis of volitional motor control. Nature Reviews Neuroscience 5:534−546.

Singh P, Jana S, Ghosal A, Murthy A. 2016. Exploration of joint redundancy but not task space variability facilitates supervised motor learning. Proceedings of the National Academy of Sciences of the United States of America 113:14414−14419.

Dimitriou M, Franklin DW, Wolpert DM. 2012. Task-dependent coordination of rapid bimanual motor responses. Journal of Neurophysiology 107:890−901.

Omrani M, Diedrichsen J, Scott SH. 2013. Rapid feedback corrections during a bimanual postural task. Journal of Neurophysiology 109:147−161.

Kobayashi T, Nozaki D. 2024. Implicit motor adaptation patterns in a redundant motor task manipulating a stick with both hands. eLife 13:RP96665.

Collins JJ. 1995. The Redundant Nature of Locomotor Optimization Laws. Journal of Biomechanics 28:251–267.

O’Sullivan I, Burdet E, Diedrichsen J. 2009. Dissociating Variability and Effort as Determinants of Coordination. PLoS Computational Biology 5:e1000345.

Selinger JC, O’Connor SM, Wong JD, Donelan JM. 2015. Humans Can Continuously Optimize Energetic Cost during Walking. Current Biology 25:2452–2456.

White O, Diedrichsen J. 2010. Responsibility assignment in redundant systems. Current Biology 20:1–6.

Zhang Z, Guo D, Huber ME, Park S-W, Sternad D. 2018. Exploiting the geometry of the solution space to reduce sensitivity to neuromotor noise. PLoS Computational Biology 14:e1006013.

Sainburg RL. 2005. Handedness: Differential Specializations for Control of Trajectory and Position. Exercise and Sport Sciences Reviews 33:206−213.

Schaffer JE, Sainburg RL. 2021. Interlimb Responses to Perturbations of Bilateral Movements are Asymmetric. Journal of Motor Behavior 53:217−233.

Yokoi A, Hirashima M, Nozaki D. 2014. Lateralized Sensitivity of Motor Memories to the Kinematics of the Opposite Arm Reveals Functional Specialization during Bimanual Actions. Journal of Neuroscience 34:9141−9151.

Johansson RS, Theorin A, Westling G, Andersson M, Ohki Y, Nyberg L. 2006. How a Lateralized Brain Supports Symmetrical Bimanual Tasks. PLoS Biology 4:e158.

Hayashi T, Yokoi A, Hirashima M, Nozaki D. 2016. Visuomotor Map Determines How Visually Guided Reaching Movements are Corrected Within and Across Trials. eNeuro 3:e0032−16.

Haggard P. 2017. Sense of agency in the human brain. Nature Reviews Neuroscience 18:197−208.

Wolpert DM, Ghahramani Z, Jordan MI. 1995. Are arm trajectories planned in kinematic or dynamic coordinates? An adaptation study. Experimental Brain Research 103:460–470.

Flanagan JR, Rao AK. 1995. Trajectory Adaptation to a Nonlinear Visuomotor Transformation: Evidence of Motion Planning in Visually Perceived Space. Journal of Neurophysiology 74:2174–2178.

Mazzoni P, Krakauer JW. 2006. An Implicit Plan Overrides an Explicit Strategy during Visuomotor Adaptation. Journal of Neuroscience 26:3642−3645.

Schaefer SY, Shelly IL, Thoroughman KA. 2012. Beside the point: motor adaptation without feedback-based error correction in task-irrelevant conditions. Journal of Neurophysiology 107:1247−1256.

Kim HE, Parvin DE, Ivry RB. 2019. The influence of task outcome on implicit motor learning. eLife 8:e39882.

Nashed JY, Crevecoeur F, Scott SH. 2012. Influence of the behavioral goal and environmental obstacles on rapid feedback responses. Journal of Neurophysiology 108:999–1009.

Oldfield RC. 1971. The Assessment and Analysis of Handedness: The Edinburgh Inventory. Neuropsychologia 9:97−113.

Scott SH. 1999. Apparatus for measuring and perturbing shoulder and elbow joint positions and torques during reaching. Journal of Neuroscience Methods 89:119−127.

Pruszynski JA, Kurtzer IL, Scott SH. 2008. Rapid Motor Responses Are Appropriately Tuned to the Metrics of a Visuospatial Task. Journal of Neurophysiology 100:224−238.

Yang L, Michaels JA, Pruszynski JA, Scott SH. 2011. Rapid motor responses quickly integrate visuospatial task constraints. Experimental Brain Research 211:231−242.

Crevecoeur F, Kurtzer I, Scott SH. 2012. Fast corrective responses are evoked by perturbations approaching the natural variability of posture and movement tasks. Journal of Neurophysiology 107:2821−2832.

